# Scale dependence decouple life history traits from transposable element evolution in sauropsids

**DOI:** 10.64898/2026.08.26.747309

**Authors:** Yun Liang, Bin Zuo, Yan-Bo Sun

**Author notes:** Correspondence: Yan-Bo Sun ( or). These authors contribute equally (Yun Liang, Bin Zuo).

## Abstract

The Mutational Hazard Hypothesis (MHH) predicts that reduced effective population size weakens purifying selection and promotes transposable element (TE) accumulation. Because effective population size is difficult to estimate across broad taxonomic scales, body mass, generation time, and *dN/dS* are often used as proxies. We analyzed TE landscapes across 167 sauropsid genomes to test whether these proxies predict genomic TE proportion consistently across phylogenetic scales. All three showed strong scale dependence. Body mass was positively associated with TE proportion across clades, but this relationship broke down within clades: only snakes retained a significant negative Pearson correlation after multiple-testing correction, whereas the corresponding phylogenetically corrected slopes were not significant. Generation time showed a strong pooled association that disappeared within every major clade in both Pearson and PGLS analyses. The pooled *dN/dS*-TE relationship also disappeared after accounting for body mass and generation time in a structural equation model, with a robust within-clade association retained only in turtles. Lineage-specific TE dynamics, including LTR expansion in sea snakes, were not captured by these proxies. These results show that commonly used MHH proxies mainly reflect clade-level structure rather than consistent within-lineage mechanisms.

**Impact statement:** Apparent links between life-history traits and transposable element accumulation across sauropsids arise largely from between-clade differences rather than consistent within-lineage mechanisms.

## Introduction

Evolutionary biology has long debated whether microevolutionary processes operating within populations can explain macroevolutionary patterns observed across deep phylogenetic timescales (Erwin, 2000; Gould, 1980; Rolland et al., 2023; Simpson, 1944). In comparative genomics, this issue often emerges when cross-species correlations between traits and genome features are interpreted as evidence for within-lineage evolutionary mechanisms. This interpretation implicitly assumes that microevolutionary processes scale up seamlessly to macroevolutionary patterns. If this assumption fails, such as when the relationship differs between the pooled cross-species level and within individual clades, the resulting inference can be misleading. This statistical manifestation of the macroevolution-microevolution gap is known as Simpson’s paradox (Simpson, 1951).

Transposable elements (TEs) evolution provides an ideal case study for examining this gap. TE proportions vary dramatically across vertebrate lineages, from low levels in most bird genomes, typically around 4-10%, to more than 50% in some non-avian reptiles (Pasquesi et al., 2018; Sotero-Caio et al., 2017). Mutational Hazard Hypothesis (MHH) (Lynch, 2007) offers an explicitly microevolutionary explanation for this variation, proposing that TE accumulation is largely a nonadaptive consequence of relaxed purifying selection in lineages with small effective population sizes (*Ne*). Because Ne is difficult to measure directly across broad taxonomic scales, comparative studies often operationalize the MHH using life-history proxies, most commonly body mass, under the assumption that larger-bodied species have smaller *Ne* (Lynch, 2007; Lynch & Conery, 2003; Marino et al., 2025). While the underlying logic operates at the population level, where variation in *Ne* should drive differences in selection efficiency against TE insertions, empirical tests typically rely on species-level averages of body mass and genome-wide TE content, creating a potential scale mismatch between the proposed mechanism and the evidence used to test it.

Empirical support for this prediction remains inconsistent across studies. At the broadest phylogenetic scale, a comparative analysis of 807 animal species found no evidence that proxies of effective population size explain long-term variation in either genome size or TE content (Marino et al., 2025). Within amniotes, the widely used selection proxy *dN/dS* shows relationships with life-history traits that differ substantially among lineages, including a particularly unexpected pattern in birds (Figuet et al., 2016). At the population level, experimental evolution in Drosophila has shown that spatially variable purifying selection can shape the genomic distribution of P-element insertions (Langmüller et al., 2023), but whether this fine-scale process scales up to explain species differences in total TE burden remains unresolved. These studies span different phylogenetic breadths, rely on different proxies for selection, and operate at different biological scales, echoing a broader problem in connecting microevolutionary processes to macroevolutionary patterns (Rolland et al., 2023). Collectively, they point to a structural problem where the relationship between selection and TE accumulation may depend on the phylogenetic scale being examined. However, comparative genomic studies rarely test whether a given predictor-TE association holds consistently within clades, or whether pooled correlations are driven mainly by between-clade differences. This issue becomes particularly acute when clades differ systematically in both life-history traits and TE content, creating conditions under which pooled correlations may arise from compositional differences rather than from repeated within-lineage evolutionary processes.

Sauropsids, comprising squamates, turtles, crocodilians, the tuatara, and birds, provide an ideal system for examining this problem directly. The major sauropsid lineages differ markedly in mean body size, generation time, and TE content (Bird et al., 2020; Mancini et al., 2025; Moura et al., 2024). Bird genomes are generally compact and TE-poor, with TEs often accounting for approximately 10% of the genome, although lineage-specific expansions have resulted in TE content reaching approximately 30% in some Piciformes (Gao et al., 2017; Manthey et al., 2018; Zhang et al., 2014). In contrast, squamate genomes show highly variable repeat content, ranging from approximately 25% to 73%, reflecting a dynamic repeat landscape (Pasquesi et al., 2018). Turtles and crocodilians also have substantial TE/repeat backgrounds (Hong et al., 2023). For instance, interspersed repeats and TEs account for approximately 38% of the genome in crocodilians (Green et al., 2014; Wan et al., 2013). Importantly, these clade-level differences in life-history traits and TE content are not randomly distributed. Birds are consistently characterized by compact genomes and relatively short generation times, whereas squamates span the widest range in both body mass and repeat content. This systematic co-variation among clade identity, life history, and TE landscape creates conditions for between-clade patterns to dominate pooled analyses, potentially masking or reversing within-lineage relationships. Furthermore, the availability of whole genomes spanning all major lineages makes it possible to conduct both pooled and within-clade analyses using a single, internally consistent dataset. This avoids the confounding effects of comparing results across studies with different taxon sampling, analytical methods, and TE annotation pipelines.

In this study, we analyzed TE landscapes across 167 sauropsid genomes to address four main questions. First, we investigated whether the relationship between body mass and TE content changes when analyses are partitioned by clade rather than pooled across all species. Second, we examined the relative contributions of generation time, body mass, and *dN/dS* to TE variation after distinguishing between-clade and within-clade components. Third, we tested whether *dN/dS*, which is often treated as a direct measure of selection efficiency, exhibits scale dependence comparable to that of life-history proxies. Finally, we used structural equation modeling (SEM) to evaluate whether *dN/dS* retains an independent association with TE content after accounting for intercorrelated life-history effects, or whether it primarily recapitulates clade-level variation. By explicitly testing for scale dependence in TE-trait associations, this study aims to provide substantive insights into sauropsid genome evolution and offer methodological cautions for future comparative genomic research.

## Results

### High-quality genomic resource and phylogenomic framework

We compiled 215 sauropsid genomes and retained 167 genomes representing all major sauropsid lineages, including Squamata, Rhynchocephalia, Testudines, Crocodylia, and Aves. Among these, 149 genomes met our primary quality criteria with BUSCO completeness at or above 85%. An additional 18 avian genomes with lower completeness (56.8% to 84.2%) were included to improve phylogenetic representation (Supplementary file 1, Table S1). We compiled species trait data including maximum body mass, generation time, and habitat from public databases and the primary literature (Supplementary file 1, Table S2). Missing generation times for 21 species were estimated using phylogenetic generalized least squares (PGLS) with body mass, climatic variables, and insularity as predictors (Supplementary file 1, Table S3), following recent generation-length modelling in amphibians and reptiles and broader evidence that body size, climate, and island occurrence covary with life-history pace (Healy et al., 2014; Mancini et al., 2025; Novosolov et al., 2013; Stark et al., 2018).

### Lineage-specific TE landscapes and phylogenetic signal

TE proportion varied substantially across the 167 sauropsid genomes, ranging from 6.21% to 54.02% (Supplementary file 1, Table S4). Mean TE proportion differed markedly among major clades, being highest in Testudines (49.05%) and lowest in Aves (10.47%). Genome size ranged from 1.30 to 4.27 Gb in non-avian reptiles and from 1.02 to 1.25 Gb in birds. Across all sampled species, TE proportion was strongly correlated with genome size (*r* = 0.76, *p* < 2.2 × 10^-16^; Figure 1—figure supplement 1), indicating that TE burden is a major contributor to genome-size variation in sauropsids.

TE composition was also strongly clade-specific (Figure 1; Figure 1—figure supplement 2). LINEs were particularly abundant in Serpentes (15.66%-28.74%), whereas DNA transposons were enriched in Testudines and Crocodylia (15.29%-26.29%). Hydrophis sea snakes additionally showed markedly elevated LTR proportion (mean 21.05%), with most LTR accumulation concentrated within the 0%-10% Kimura-divergence range (Figure 1—figure supplement 3). In contrast, birds exhibited low total TE proportion (mean 10.47%) and generally streamlined repeat landscapes. Similar lineage-specific and dynamic repeat landscapes have been reported in snake genomes and across squamate reptiles (Castoe et al., 2011; Pasquesi et al., 2018).

**Figure 1.**
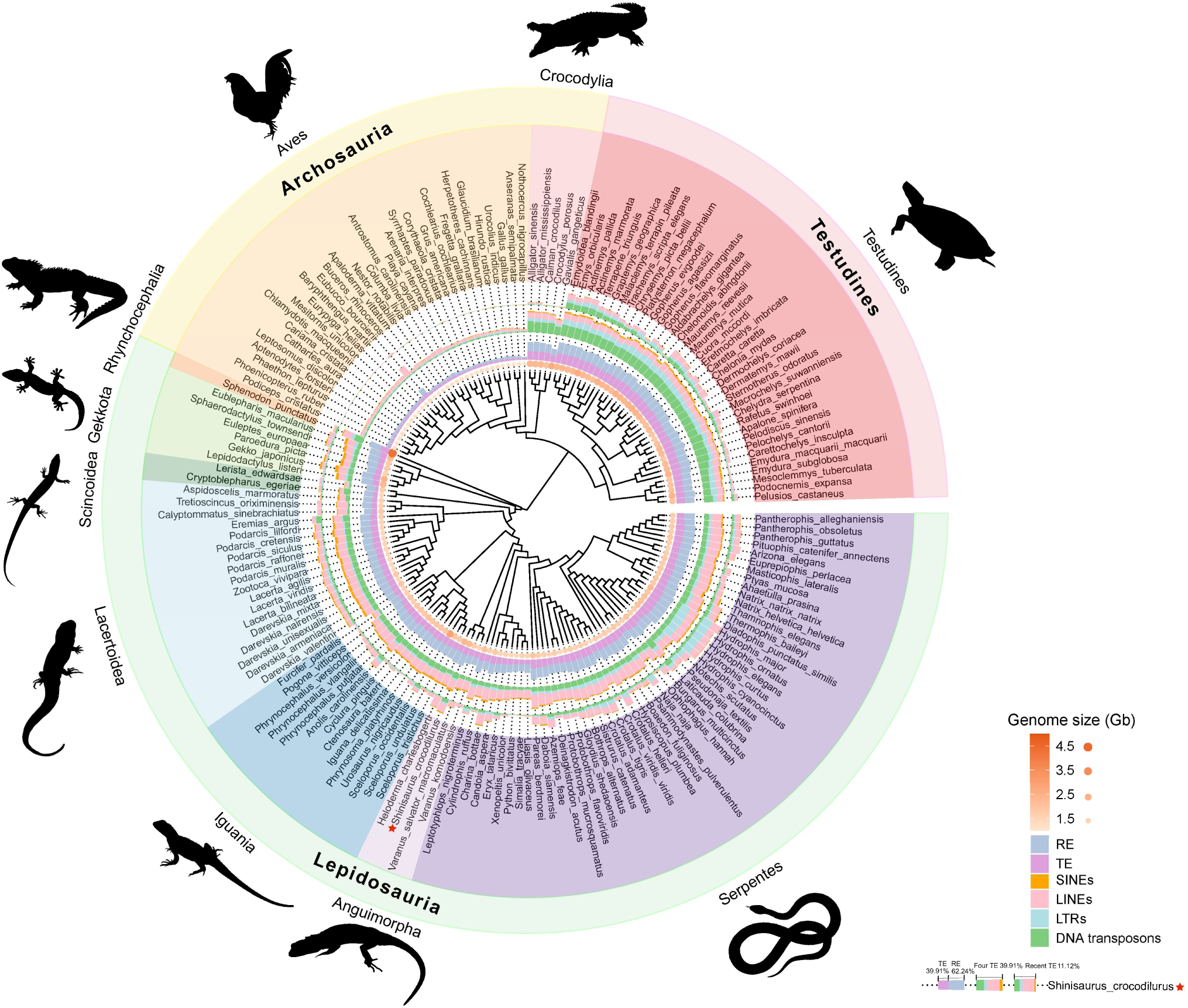
Phylogenetic tree and TE dynamics across reptilian lineages. From inner to outer rings: (1) The bubble plot illustrates genome size variation, where larger and darker-colored bubbles indicate greater genome sizes. (2) Stacked bar chart delineates repetitive sequence and TE proportions. (3) Secondary stacked bar chart breaks down TE class composition. (4) Tertiary stacked bar chart specifies TE class distribution among recent insertions (divergence ≤ 5%). SINEs, LINEs, LTRs, and DNA transposons are classified as major TE classes.

Low-divergence TE copies with Kimura divergence ≤5%, used here as an approximate indicator of recent TE accumulation (Palacios-Gimenez et al., 2020), similarly showed pronounced lineage-specific patterns (Figure 1; Supplementary file 1, Table S5). Recent LINE accumulation was prominent in squamates (average 3.28%). Hydrophis species showed particularly high levels of both low-divergence LINEs (mean 5.96%) and LTRs (mean 6.86%). In stark contrast, Crocodylia, Pythonidae, and birds contained relatively few low-divergence TE copies. This paucity is consistent with lineage-specific genomic contexts reported previously, including exceptionally slow genome evolution in crocodilians (Green et al., 2014), low identifiable repeat content in python relative to some other snakes (Castoe et al., 2011), and efficient DNA removal contributing to compact avian genomes (Kapusta et al., 2017). More generally, lineage-specific differences in host defence, including piRNA-mediated TE silencing, may also contribute to variation in recent TE activity, although this mechanism was not directly tested here (Ozata et al., 2019).

All major TE classes exhibited strong phylogenetic signal (Pagel’s λ = 0.95-0.99; Supplementary file 1, Table S6). Ancestral state reconstruction similarly identified distinct lineage-specific trajectories, including increased LINE content in snakes, increased LTR content in marine snakes and turtles, and reductions across major TE classes in birds (Supplementary file 1, Table S7; Figure 1—figure supplements 4-8). These results demonstrate that variation in TE abundance and composition is deeply structured by shared evolutionary history, motivating explicit separation of between-clade and within-clade relationships in subsequent analyses.

### Scale-dependent reversal of the body mass and TE relationship

A cross-clade correlation analysis across 167 sauropsid species revealed a positive relationship between body mass and TE proportion (*r* = 0.213, *p* = 0.0056; Figure 2a and Supplementary file 1, Table S8). This initial result suggested that larger species might carry a higher TE proportion. However, this apparent trend actually stems from pronounced compositional disparities among major clades. Testudines and Crocodylia are not only massive in size (mean body mass 52.83 kg and 472.64 kg, respectively) but also possess high TE proportion (mean TE 49.05% and 43.18%, respectively), whereas birds are small in size (2.17 kg) and possess low TE proportion (mean TE 10.47%). When the analysis was restricted to individual clades, the pooled positive relationship broke down, with a robust negative correlation in Serpentes and nominal negative trends in Aves, a pattern consistent with Simpson’s paradox rather than evidence for a universal within-clade relationship (Figure 2c and Supplementary file 1, Tables S9 and S10).

**Figure 2.**
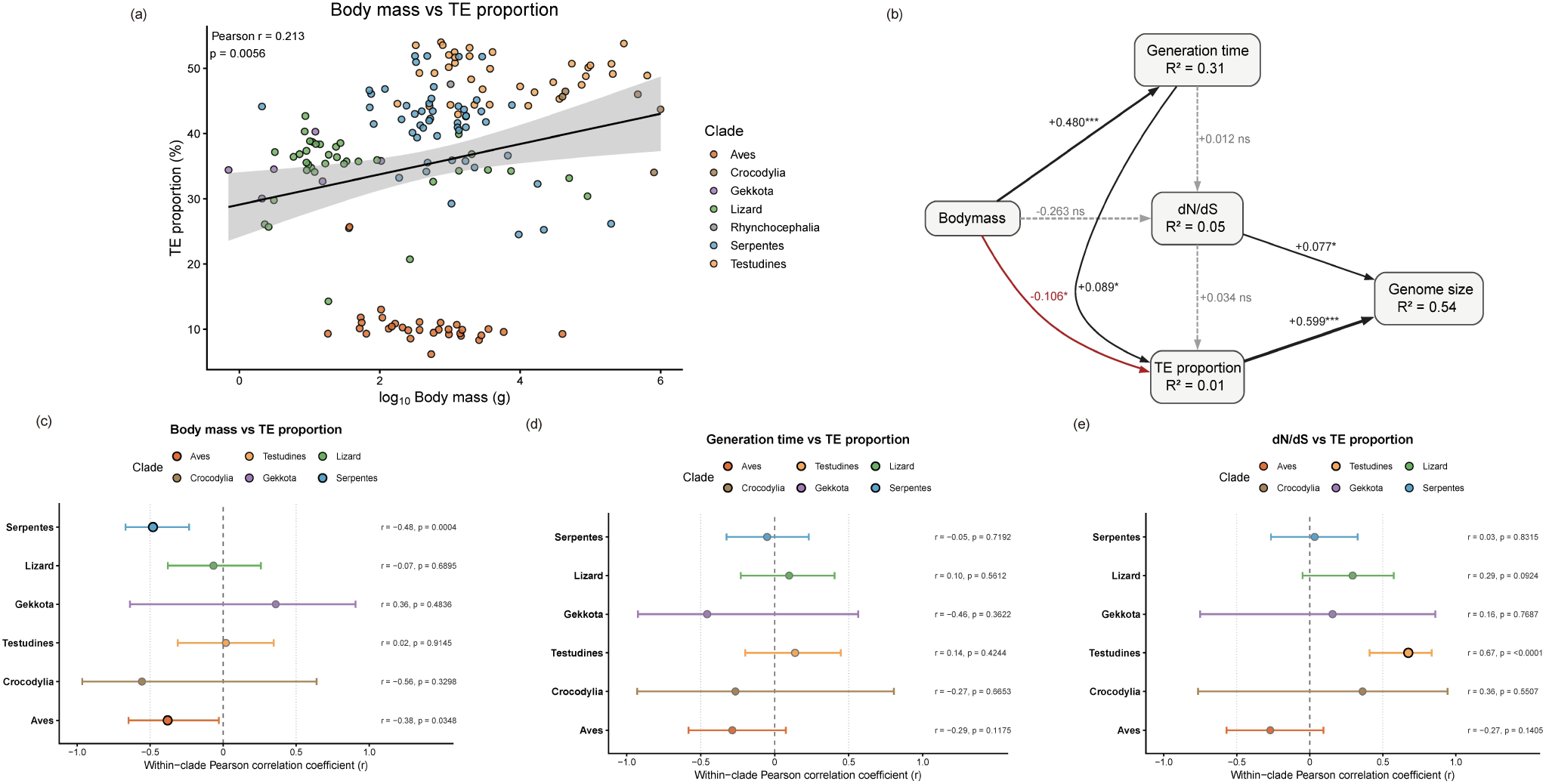
A Simpson’s Paradox in sauropsid TE evolution. (a) Cross-clade scatter plot of log10-transformed body mass versus TE proportion for all 167 sauropsids, colored by clade. The pooled Pearson correlation is positive (*r* = 0.213). (b) Results of the segmented structural equation model. Fisher’s C = 2.715, *p* = 0.607; Chi-squared = 1.158, *p* = 0.56. The black solid line represents a positive correlation, the red solid line represents a negative correlation, the gray line represents no correlation, and the thickness of the line represents the degree of correlation. TE proportion and *dN/dS* jointly explain 54% of the variation in genome size (R² = 0.54). Weight explains 31% of the variation in generation time (R² = 0.31). The explanatory degree of *dN/dS* is (R² = 0.05), and the explanatory degree of TE proportion is (R² = 0.01). (c) Forest plot presenting the results of person-related correlation analyses of body mass and TE proportion in each group. (d) Forest plot presenting the results of person-related correlation analyses of generation time and TE proportion in each group. (e) Forest plot presenting the results of person-related correlation analyses of *dN/dS* and TE proportion in each group.

This reversal was particularly pronounced within Serpentes (n = 50), where body mass showed a strong negative correlation with TE proportion (*r* = -0.481, nominal *p* = 0.0004, BH-adjusted *p* = 0.0025). The corresponding PGLS slope was also negative (*β* = -0.0169, nominal *p* = 0.0490), but it did not remain significant after BH correction (BH-adjusted *p* = 0.1469). Importantly, this negative within-clade pattern is unlikely to be explained solely by genome-size fluctuation. For example, the 197.7 kg *Python bivittatus* contains only 26.19% TE, whereas the diminutive 70.8 g *Pareas berdmorei* snake possesses a TE proportion as high as 46.63%. Despite a 2,790-fold difference in body mass, their overall genome sizes are remarkably similar (∼1.44 Gb vs. ∼1.85 Gb, respectively), meaning the absolute TE payload in *P. berdmorei* is nearly twice that of the giant python. Consistent with this, univariate PGLS models incorporating absolute TE amount showed that body mass fails to provide a stable, independent positive predictor for absolute TE burden across sauropsids (coefficient = -0.0164, *p* = 0.0858; conditionally negative when controlling for genome size, coefficient = -0.0107, *p* = 0.0364) (Supplementary file 1, Table S11).

Similar negative or null within-clade patterns exist in other clades. Birds (n = 31) showed negative nominal associations in both analyses (Pearson *r* = -0.380, nominal *p* = 0.0348; PGLS *β* = -0.0378, nominal *p* = 0.0226), but neither association remained significant after BH correction (Pearson BH-adjusted *p* = 0.1044; PGLS BH-adjusted *p* = 0.1355). Crocodilians (n = 5) also displayed a negative trend (*r = -*0.557*, p =* 0.330; PGLS *β* = -0.046, *p* = 0.360), though it was statistically insignificant due to the small sample size. Turtles, lizards and geckos showed no significant correlation. The significant reversal in Serpentes, together with the disappearance of the pooled positive relationship in the remaining clades, is consistent with a Simpson’s paradox-like aggregation effect (Simpson, 1951). This indicates that the observed positive cross-clade correlation does not reflect a universal within-clade mechanism directly linking large body size to TE accumulation. Rather, it reflects a scale-dependent aggregation effect arising from phylogenetic structure and compositional differences among clades.

### Generation time tracks cross-clade TE variation but shows signal collapse within clades

We next examined generation time, a core life-history trait that is often assumed to relate more directly to TE dynamics because it influences the number of reproductive cycles per unit evolutionary time. Across 167 sauropsid species, generation time was positively correlated with TE proportion (*r* = 0.517, *p <* 0.0001) (Supplementary file 1, Table S8 and Figure 2—figure supplement 1). When major clades were ranked by mean TE proportion, the ranking broadly tracked their average generation time: Testudines (29.04 years; 49.05% TE) > Crocodylia (23.0 years; 43.18% TE) > Serpentes (9.08 years; 41.25% TE) > Gekkota (5.03 years, 34.64% TE) > Lizards (5.9 years, 34.35% TE) > Aves (6.28 years; 10.47% TE). Although this macroevolutionary gradient was not perfectly monotonic for generation time, it clearly indicates that generation time captures a broad cross-clade macroevolutionary signal of TE accumulation. The most notable deviation from this trend occurs in birds, which maintain highly streamlined genomes and atypically low TE burdens despite having generation times comparable to those of small squamates.

To further evaluate the relative contributions of generation time, body mass, and *dN/dS* to TE accumulation, we used piecewise structural equation modeling (SEM; Figure 2b and Table 1). In the SEM, generation time remained significantly positively associated with TE proportion (*β* = 0.089, *p* = 0.024), indicating that this cross-clade signal persisted after accounting for body mass and *dN/dS*. Body mass showed a significant negative direct effect on TE proportion (*β* = -0.106, *p* = 0.015), while also being positively associated with generation time (*β* = 0.480, *p* < 0.001). In contrast, *dN/dS* did not significantly predict TE proportion in this model (*β* = 0.034, *p* = 0.180).

**Table 1.**
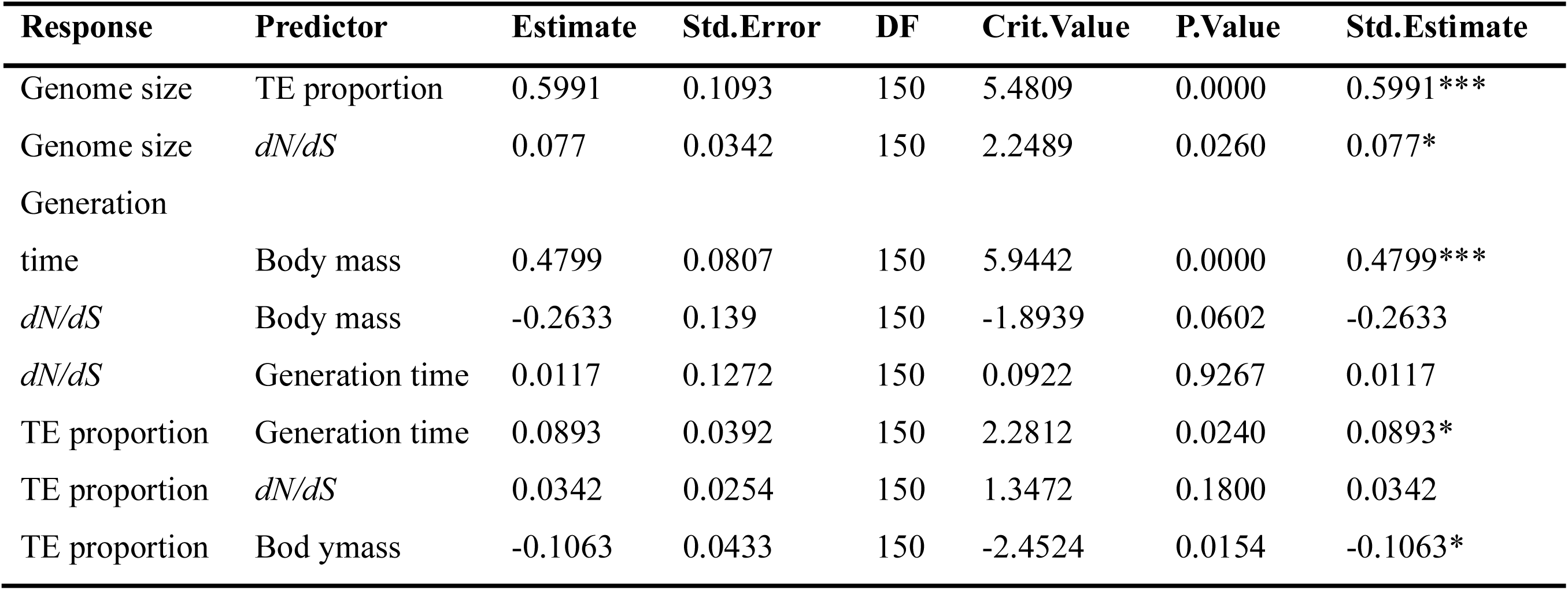
Structural equation model results. Overall goodness-of-fit: Fisher’s C = 2.715 (*p* = 0.607), Chi-squared = 1.158 (*p* = 0.56), meeting the goodness-of-fit criterion (p > 0.05). The model explained 54% of the variance in genome size, 31% of the variance in generation time, and smaller amounts of variance: 5% for *dN/dS* and 1% for TE proportion. All coefficients have been phylogenetically corrected. Significance codes: 0 ‘***’ 0.001 ‘**’ 0.01 ‘*’ 0.05.

However, analyses based on absolute TE amounts revealed that this cross-clade signal is largely indirect and deeply intertwined with genome-size dynamics. In our pooled absolute-TE PGLS models, log10 generation time alone did not significantly predict log10 absolute TE amount (coefficient = 0.0324, *p* = 0.311). When log10 genome size was included as a covariate, the generation-time coefficient was further reduced and remained non-significant (coefficient = 0.00583, *p* = 0.714) (Supplementary file 1, Table S11). These results indicate that generation time does not independently explain absolute TE amount once genome-size variation and phylogenetic non-independence are taken into account.

Furthermore, the generation time signal proved strongly scale-dependent. When generation time and TE proportion were analyzed separately within each major clade, the positive cross-clade signal was not consistently retained. Aves showed nominally significant negative associations in both the Pearson analysis (*r* = -0.383, nominal *p* = 0.0336) and PGLS (*β* = -0.295, nominal *p* = 0.0328), but neither remained significant after BH correction (BH-adjusted *p* = 0.2016 and 0.1966, respectively). No major clade retained a significant generation-time association after multiple-testing correction (Figure 2d and Supplementary file 1, Tables S9 and S10). Unlike the body mass-TE relationship, this pattern does not represent Simpson’s paradox, because there was no consistent reversal in direction between pooled and within-clade analyses. Instead, it represents a form of signal collapse: a strong cross-clade association disappears entirely at the within-clade scale. Thus, generation time does not drive an independent, universal microevolutionary accumulation of TEs; rather, it serves as a macroevolutionary correlate of broader covariation among life history, genome size, and deep-time lineage divergence.

### Clade-specific heterogeneity in the *dN/dS* and TE relationship

Although *dN/dS* did not independently predict TE proportion in the SEM, it is commonly used as a molecular proxy for the efficacy of purifying selection. We therefore further examined whether the relationship between *dN/dS* and TE proportion remained consistent across phylogenetic scales. Using a matched dataset of 150 species (FitMG94-derived), we recovered a strong positive pooled cross-clade correlation between TE proportion and *dN/dS* (*r* = 0.616, *p* < 0.0001; n = 150; Supplementary file 1, Table S8 and Figure 2—figure supplement 2), superficially supporting the expectation that lineages under weaker purifying selection accumulate more TEs.

However, much like body mass and generation time, this proxy exhibited strong scale dependence and was closely entangled with genome-size dynamics (Figure 2e and Supplementary file 1, Tables S9 and S10). Within-clade analyses revealed strong clade-specific heterogeneity. Only Testudines retained a significant association after multiple-testing correction (Pearson *r* = 0.674, nominal *p* = 6.09 × 10⁻⁵, BH-adjusted *p* = 0.0004; PGLS *β* = 0.804, nominal *p* = 0.0016, BH-adjusted *p* = 0.0095; n = 29). No significant association was retained in Aves, Serpentes, Lizards, Crocodylia, or Gekkota after BH correction.

When transitioning to absolute TE amount, a univariate PGLS initially showed a positive association with *dN/dS* (coefficient = 0.408, *p* = 0.0357). However, introducing log10 genome size as a covariate substantially reduced the *dN/dS* coefficient, but did not eliminate the association (coefficient = 0.0747, *p* = 9.54 × 10⁻⁵) (Supplementary file 1, Table S11). These results indicate that the pooled *dN/dS*-TE relationship is strongly shaped by genome-size dynamics, although a weak residual positive association with *dN/dS* remains.

Taken together, these integrated analyses demonstrate that cross-clade correlations between biological proxies (body mass, generation time, *dN/dS*) and TE burden are confounded by phylogenetic structure and genome-size variation. None of these proxies provides a universal, clade-consistent explanation for TE accumulation across sauropsids.

## Discussion

Analyses of 167 sauropsid genomes show that pooled cross-clade TE correlations do not necessarily hold within major clades. Cross-clade correlations can reveal broad macroevolutionary patterns, but they do not automatically represent within-clade evolutionary mechanisms. Body mass, generation time, and *dN/dS* therefore appear to capture broad among-clade differences in TE proportion, but they do not provide scale-consistent predictors of within-clade TE variation in Sauropsida. Crucially, these life-history traits often covary with genome size baselines at the macroevolutionary scale, masking lineage-specific dynamics of TE accumulation. Therefore, research should shift focus from broad life-history associations to the actual lineage-specific mechanisms driving intra-clade TE variation.

### Life-history proxies show scale-dependent and non-independent effects

Our results indicate that body mass and generation time are positively correlated with TE proportion at the whole-dataset scale. Larger-bodied species tend to exhibit longer generation times, lower fecundity, and delayed sexual maturity. Thus, at the inter-clade scale, body mass and generation time may jointly reflect a macroevolutionary background often associated with reduced long-term effective population size (*Ne*) and an increased probability of TE retention. According to the generation time hypothesis (Lewin et al., 2025; Thomas et al., 2010), species with shorter generations typically undergo more germline replication cycles per unit time, potentially increasing replication-associated mutations. However, TE load is not merely a product of the *de novo* insertion rate. It represents the net outcome of new insertions, deletions, epigenetic silencing (Almeida et al., 2022), and purifying selection (Langmüller et al., 2023). Consequently, the positive correlation observed at the inter-clade scale is more appropriately interpreted as a macroevolutionary baseline of long-term TE retention rather than a direct mechanistic driver.

Crucially, this inter-clade trend collapses at the intra-clade level. Within specific clades, the opposing forces of generation time likely counteract each other. While longer generations reduce replication cycles and novel insertions, they also associate with smaller *Ne* and weaker purifying selection. Furthermore, intra-clade TE variation is more likely driven by TE family bursts at specific nodes or horizontal transfers within specific lineages rather than gradual life-history shifts. For example, Copia and Gypsy LTR elements in *Hydrophis curtus* expanded independently at approximately 7 Ma and 12 Ma, respectively (Peng et al., 2020), while *Aipysurus laevis* experienced multiple horizontal transfer events of LINEs following its marine transition (Galbraith et al., 2020).

The scale dependency was most pronounced for body mass, with negative estimates concentrated in snakes and birds. However, after BH correction, only the Pearson association in snakes remained significant; the bird associations and all within-clade body-mass PGLS slopes were not statistically significant. In snakes, previous studies have revealed highly differentiated repetitive landscapes where larger-bodied (e.g. Pythonidae) paradoxically contain fewer identifiable repeats (Ahmad et al., 2021). This indicates that divergent histories of TE amplification and deletion rates override the effects of body mass. In birds, the nominal negative trend, which did not remain significant after BH correction, may be linked to genomic constraints associated with flight. The high metabolic rates and small cell sizes associated with powered flight are thought to constrain genome size, thereby limiting repeat expansion and shaping avian retrotransposon composition (Ji & DeWoody, 2016). Therefore, the lower TE proportions in larger-bodied birds likely reflect an integrated suite of physiological and genome size constraints rather than a simple body mass effect.

### Limits of *dN/dS* as a proxy for TE accumulation

The *dN/dS* ratio is widely used as a molecular proxy for the efficacy of purifying selection. While FitMG94-derived *dN/dS* showed a significant positive correlation with TE proportion across the entire dataset (*r* = 0.616, *p* < 0.0001), superficially supporting the Mutational Hazard Hypothesis (MHH), this relationship weakened under multivariate and scale-explicit scrutiny. In the structural equation model (SEM), *dN/dS* failed to independently predict TE proportion after accounting for body mass and generation time. Furthermore, within-clade analyses showed that the TE proportion-*dN/dS* relationship remained significant after BH correction only in turtles in both Pearson and PGLS analyses, whereas no association was retained in the other major clades. These results indicate that *dN/dS* is not a universal, clade-consistent predictor of TE proportion across sauropsids.

Existing theoretical frameworks suggest that reduced *Ne* compromises the selective removal of weakly deleterious TE insertions (Bourgeois & Boissinot, 2019; Lynch & Conery, 2003; Mérel et al., 2025). However, large-scale comparative studies increasingly suggest that Ne alone does not stably explain long-term variation in genome size or TE content (Marino et al., 2025). For instance, according to the nearly neutral theory, larger-bodied, longer-lived birds should possess smaller *Ne* and experience reduced purifying selection, which would manifest as elevated *dN/dS* values; yet, this expected trend is not observed (Figuet et al., 2016). This highlights that the theoretical link between life-history-mediated selection efficacy and *dN/dS* is not universally conserved across deeply diverged amniote lineages.

The results from crocodilians further underscore the limitations of using *dN/dS* as a standalone predictor for TE accumulation. Despite sharing trait profiles typically associated with relaxed selective constraints, such as long generation times, slow life histories, and high overall TE content, crocodilians exhibit no significant *dN/dS*-TE association within their clade. While this null result may be partly attributable to limited sample sizes and a narrow range of intra-clade variance, a plausible biological explanation lies in the exceptionally slow genomic evolution of crocodilians. Genomic studies have highlighted that crocodilian exhibit exceptionally slow evolutionary rates across multiple dimensions, including nucleotide substitutions, indels, TE content and transposition activity, gene family evolution, and chromosomal synteny (Green et al., 2014). Consequently, this broad reduction in evolutionary rate may limit detectable intra-clade covariation between *dN/dS* and TE proportion.

In summary, the apparent positive cross-clade correlation between *dN/dS* and TE proportion is best interpreted as a scale-dependent macroevolutionary association rather than direct evidence for a universal mechanism of relaxed purifying selection driving TE accumulation. Conversely, the association that remained significant after multiple-testing correction uniquely in turtles more likely represents a case of historical covariation rather than a ubiquitous mechanistic rule.

### Methodological Insights in Comparative Genomics

Previous comparative studies have typically employed global PGLS models or pooled correlations to examine the TE-trait relationships (Felsenstein, 1985; Ji et al., 2022; Lartillot & Poujol, 2010; Marino et al., 2025; Mérel et al., 2025; Szitenberg et al., 2016; Wu & Lu, 2019). While these methods provide an important framework for identifying macroevolutionary patterns, a critical issue must be explicitly acknowledged: cross-clade correlations do not necessarily equate to within-clade mechanisms. Although PGLS corrects for phylogenetic non-independence among species caused by shared ancestry, a single global PGLS model still estimates an average slope across the sampled phylogeny and may obscure heterogeneity among major lineages. Our within-clade PGLS estimates were often directionally consistent with the Pearson correlations, but the statistical support was substantially weaker after multiple-testing correction. No body-mass or generation-time PGLS association remained significant after BH correction, whereas the *dN/dS* association was retained only in Testudines, indicating that phylogenetic correction alone does not restore a universal relationship between MHH-related proxies and TE proportion. Together, these results reinforce the absence of scale-consistent relationships across clades, while identifying a single robust clade-specific *dN/dS* signal in Testudines. Relying solely on global correlations or single-slope PGLS models therefore risks interpreting historical baseline differences among clades as evidence for universal mechanisms.

Future comparative genomic studies must prioritize slope heterogeneity and scale consistency. Researchers should routinely report within-clade slopes and explicitly test for predictor-by-clade interactions. More importantly, future studies should not rely on TE proportion alone. Because TE proportion is a ratio determined jointly by absolute TE length and total genome size, it is inevitably coupled to genome-size dynamics. Comparative genomic analyses should therefore also incorporate absolute TE abundance in base pairs or megabases and treat total genome size as a key genomic constraint. Finally, distinguishing between long-term retention and recent expansion requires high-resolution metrics such as TE age structure (Kimura divergence) and family-specific analyses, shifting the field’s focus toward the actual epigenetic and genomic defense mechanisms that shape lineage-specific evolution.

## Materials and Methods

### Data collection and genome completeness assessment

Genome assemblies of sauropsids were obtained from the National Center for Biotechnology Information (NCBI), the National Genomics Data Center (NGDC), and the Bird Genome 10K Project database (B10K). The dataset spans the major sauropsid lineages, including Squamata (lizards, snakes, and geckos), Rhynchocephalia (Sphenodon punctatus), Testudines (turtles), Crocodylia (crocodilians), and Aves. A total of 215 candidate genome assemblies were initially collected. Genome quality was evaluated using two complementary metrics: (1) completeness, assessed using BUSCO v5.5.0 (Seppey et al., 2019) with the vertebrata_odb10 dataset (Kriventseva et al., 2019), which comprises 3,354 conserved vertebrate single-copy orthologs; and (2) assembly continuity, measured by scaffold N50 and contig N50 statistics. Assemblies with BUSCO completeness ≥85% were generally retained, and this threshold was met by 149 species. To preserve broad phylogenetic representation of Aves, an additional 18 avian assemblies with BUSCO completeness of 56.8-84.2% were retained and explicitly identified in Supplementary File 1, Table S1. Scaffold N50 and contig N50 were used as descriptive measures of assembly continuity rather than as independent exclusion criteria. The final dataset comprised 167 sauropsid species for downstream analyses.

Species specific trait data were compiled from several authoritative databases and literature sources. These data were obtained from the Reptile Database (Uetz, 2024), Animal Diversity Web (Myers, 2024), the IUCN Red List, and peer-reviewed studies (Bird et al., 2020; Moura et al., 2024; Oskyrko et al., 2024). For 21 species lacking reported generation time, values were estimated using phylogenetic generalized least squares (PGLS) with body mass, climatic factors, and insularity as predictors, following the methodology of Mancini (Mancini et al., 2025). The collected datasets are available in Supplementary File 1, Table S2.

### Transposable element annotation

For each of the 167 genomes, TEs were annotated using a combined de novo and homology-based pipeline. De novo species-specific consensus repeat libraries were identified using RepeatModeler v2.0.5 (Flynn et al., 2020) with default parameters. Subsequently, the library was merged with Repbase v2018 (Bao et al., 2015) and Dfam v3.8 (Storer et al., 2021) databases to create a comprehensive reference for TE classification. Further, RepeatMasker v4.1.5 (Tarailo-Graovac & Chen, 2009) was used to annotate TEs in 167 genomes. TEs were classified into four major classes: SINEs (short interspersed nuclear elements), LINEs (long interspersed nuclear elements), LTRs (long terminal repeats), and DNA transposons. TE copies with Kimura divergence ≤5% were treated as an approximate indicator of recent TE accumulation.

### Phylogenetic inference and *dN/dS* calculation

For phylogenetic reconstruction, OrthoFinder v3.0.1b1 (Emms & Kelly, 2015) was implemented with the -S diamond_ultra_sens parameter to cluster protein sequences from 167 species. Given that merely 36 single-copy orthologs were detected across all taxa, we selected 1,193 orthogroups that were represented by single-copy genes in at least 160 of the 167 species (≥95.8% taxon occupancy) to enhance phylogenetic resolution and accuracy. The concatenated alignment of these 1,193 orthogroups was subsequently employed to reconstruct the species tree using FastTree v2.1.11 (Price et al., 2010).

To assess selection pressure on protein-coding genes, the longest transcript of each species was extracted based on 1,193 orthogroups. Protein sequences were aligned using MAFFT v7.525 (Katoh & Standley, 2013), and the resulting amino acid alignments were converted back to codon alignments with PAL2NAL v14.1 (Suyama et al., 2006). After excluding low-quality genes based on gap-free criteria, 1,047 high-quality orthologous genes were retained for subsequent analyses. Branch-specific *dN/dS* ratios (ω) were estimated using the FitMG94 model implemented in HyPhy v2.5.94 (Kosakovsky Pond et al., 2020). To mitigate biases arising from alignment errors, taxon misassignment, or unstable parameter estimates (Albà & Castresana, 2005; Zuo et al., 2024), we applied stringent filters: only values within the biologically interpretable range (0 < ω < 1) were retained, while non-positive values or values ≥1 were censored. Subsequently, branches with fewer than 155 valid ω estimates, corresponding to the first quartile (Q1) of valid gene counts across branches, were excluded to enhance statistical power. For each species, genome-wide *dN/dS* was calculated as the arithmetic mean of valid gene-specific ω estimates from the corresponding terminal branch. The resulting FitMG94-derived estimates were used as the primary metric of background selective pressure in downstream comparative analyses (Figure 2—figure supplement 3 and Supplementary file 1, Table S12).

### Ancestral State Reconstruction and Phylogenetic Signal

Ancestral states of TE proportions (SINEs, LINEs, LTRs, and DNA transposons) were reconstructed based on the percentage of the genome occupied by each TE category using maximum-likelihood estimation implemented with the fastAnc function in phytools v2.3.0 (Revell, 2011) in R v4.5.2. To evaluate phylogenetic signals in major TE categories, Pagel’s (λ) was estimated for the genomic proportion of each TE category using the phylosig function in phytools, with (λ=1) indicating a phylogenetic covariance structure consistent with Brownian-motion evolution and (λ=0) indicating no phylogenetic signal. Evolutionary patterns were visualized using the contMap function, which generated phylogeny-anchored color-gradient maps showing lineage-specific variation in TE genomic proportions.

### Cross-scale analyses of MHH-related predictors of TE proportion

To test whether commonly used Mutational Hazard Hypothesis (MHH)-related proxies provide scale-consistent predictors of genomic TE proportion, we analyzed the relationships between TE proportion and three predictors: body mass, generation time, and genome-wide *dN/dS*. TE abundance was analyzed as genomic TE proportion and expressed as percentage values in the Pearson correlation analyses. Body mass and generation time were obtained from the species-level trait table, whereas *dN/dS* values were obtained from FitMG94-derived branch-specific estimates. For *dN/dS* analyses, only species with both TE proportion and *dN/dS* estimates were retained.

We first performed cross-scale Pearson correlation analyses to compare the direction and strength of trait-TE relationships between analytical scales. Analyses were conducted at two scales: the pooled cross-clade scale, using all available sauropsid species, and the within-clade scale, using major sauropsid clades separately. Within-clade analyses were performed for Serpentes, Lizard, Gekkota, Testudines, Crocodylia, and Aves. Rhynchocephalia was retained in summary tables but excluded from within-clade interpretation because it was represented by a single species. Body mass and generation time were log10-transformed in both pooled and within-clade analyses, whereas *dN/dS* was analyzed without log transformation. For each predictor, within-clade p-values were adjusted across the six major clades using the Benjamini-Hochberg procedure. Both nominal and adjusted p-values were reported, and adjusted p-values were used for inference.

To account for phylogenetic non-independence, we further performed within-clade phylogenetic generalized least-squares analyses as phylogenetically corrected robustness tests. PGLS models were fitted separately for clades containing at least five species, including Serpentes, Lizard, Gekkota, Testudines, Crocodylia, and Aves. Models were implemented using caper v1.0.4 (Orme et al., 2011) in R v4.5.2, with Pagel’s λ estimated by maximum likelihood. TE proportion was natural-log-transformed, body mass was log-transformed, generation time was log10-transformed, and *dN/dS* was analyzed without transformation. The focal predictor p-values were adjusted across clades using the same Benjamini-Hochberg procedure.

To determine whether associations based on TE proportion were influenced by genome-size variation, we also performed pooled PGLS analyses using absolute TE amount as the response variable. Absolute TE amount was calculated as the summed length of SINEs, LINEs, LTRs, and DNA transposons. Absolute TE amount, genome size, body mass, and generation time were log10-transformed, whereas *dN/dS* remained untransformed. For each predictor, models were fitted with and without genome size as a covariate.

Finally, we used a phylogenetically informed piecewise structural equation model, implemented using the piecewiseSEM v2.3.1 (Lefcheck, 2016) in R v4.5.2, to evaluate the joint relationships among body mass, generation time, *dN/dS*, TE proportion, and genome size. Model adequacy was evaluated using Fisher’s C statistic and tests of directed separation. Body mass, generation time, and genome size were log10-transformed, and all variables were standardized before analysis. Component models were fitted using generalized least squares with a Brownian phylogenetic correlation structure. The SEM included 150 species with complete *dN/dS* estimates. This combined framework allowed us to distinguish three cross-scale patterns: sign reversal between pooled and within-clade relationships, interpreted as Simpson’s paradox; disappearance of a pooled association within clades, interpreted as signal collapse; and clade-restricted significance, interpreted as clade-specific heterogeneity.

## Acknowledgments

This work was supported by National Key Research Development Program of China (2022YFF0802300) and Yunnan Fundamental Research Projects (202401BC070011).

## Author Contributions

Yan-Bo Sun conceived the research; Yun Liang and Bin Zuo collected all the data and conducted the analyses; Yun Liang, Bin Zuo, and Yan-Bo Sun wrote and revised the manuscript.

## Data Availability

The newly generated gene structure annotations for 130 species, functional annotations for 167 species, whole-genome alignment MAF files for 138 species, and the associated analysis code have been deposited in Zenodo (https://zenodo.org/records/17301220).

## Figure supplement legends

**Figure 1—figure supplement 1. Relationship between genome size and transposable element proportion across sauropsids.**

Genome size was strongly positively correlated with genomic TE proportion across the sampled sauropsids (Pearson’s *r* = 0.76, p < 2.2×10^−16^). The solid line represents the fitted linear relationship, and the shaded area represents the 95% confidence interval.

**Figure 1—figure supplement 2. Variation in major transposable element classes among sauropsid taxonomic groups.**

Boxplots show the genomic proportions of (A) SINEs, (B) LINEs, (C) LTRs, and (D) DNA transposons across major taxonomic groups. Individual points represent species. Anguimorpha, Iguania, Lacertoidea, and Scincoidea were combined into the operational “Lizard” category.

**Figure 1—figure supplement 3. Transposable element divergence landscapes in five sea snake species.**

Divergence landscapes are shown for *Hydrophis elegans*, *Hydrophis curtus*, *Hydrophis cyanocinctus*, *Hydrophis major*, and *Hydrophis ornatus*. Bars represent the genomic proportions of SINEs, LINEs, LTRs, and DNA transposons across Kimura-divergence intervals. Most LTR accumulation was concentrated within the 0-10% divergence interval.

**Figure 1—figure supplement 4. Ancestral-node numbering in the sauropsid phylogeny.**

Numbers shown at internal nodes identify the ancestral branches used for ancestral-state reconstruction. Node 168 represents the most recent common ancestor of the 167 species included in this study.

**Figure 1—figure supplement 5. Ancestral-state reconstruction of SINE proportion across sauropsids.**

Branch colors represent reconstructed ancestral and observed tip values for the genomic proportion of SINEs according to the continuous color scale.

**Figure 1—figure supplement 6. Ancestral-state reconstruction of LINE proportion across sauropsids.**

Branch colors represent reconstructed ancestral and observed tip values for the genomic proportion of LINEs according to the continuous color scale.

**Figure 1—figure supplement 7. Ancestral-state reconstruction of LTR proportion across sauropsids.**

Branch colors represent reconstructed ancestral and observed tip values for the genomic proportion of LTRs according to the continuous color scale.

**Figure 1—figure supplement 8. Ancestral-state reconstruction of DNA transposon proportion across sauropsids.**

Branch colors represent reconstructed ancestral and observed tip values for the genomic proportion of DNA transposons according to the continuous color scale.

**Figure 2—figure supplement 1. Cross-clade relationship between generation time and transposable element proportion.**

Scatter plot showing the relationship between log10-transformed generation time and genomic TE proportion across all 167 sauropsids. Points are colored by major taxonomic group. The solid line represents the fitted linear relationship, and the shaded area represents the 95% confidence interval.

**Figure 2—figure supplement 2. Cross-clade relationship between *dN/dS* and transposable element proportion.**

Scatter plot showing the relationship between branch-level *dN/dS* and genomic TE proportion across the species retained after evolutionary-rate filtering. Points are colored by major taxonomic group. The solid line represents the fitted linear relationship, and the shaded area represents the 95% confidence interval.

**Figure 2—figure supplement 3. Distribution of gene coverage across phylogenetic branches and determination of the filtering threshold.**

The histogram shows the number of phylogenetic branches with different numbers of genes yielding valid *dN/dS* estimates. The red dashed line indicates the first quartile of the distribution (Q1=155). Only branches represented by at least 155 genes and satisfying the specified *dN/dS* filtering criteria were retained for downstream analyses.

## Supplementary files

**Supplementary File 1. Supplementary Tables S1-S12.**

**Supplementary Table S1.** BUSCO assessment results.

**Supplementary Table S2.** Trait information and data sources.

**Supplementary Table S3.** Predicted generation times for 21 species.

**Supplementary Table S4.** Genome-wide transposable element annotation results.

**Supplementary Table S5.** Transposable elements with Kimura divergence ≤ 5%

**Supplementary Table S6.** Phylogenetic signal estimates for major transposable element classes.

**Supplementary Table S7.** Ancestral-state reconstruction data for major transposable element classes.

**Supplementary Table S8.** Cross-clade correlation analyses.

**Supplementary Table S9.** Within-clade correlation analyses.

**Supplementary Table S10.** Phylogenetic generalized least-squares analyses of transposable element proportion.

**Supplementary Table S11.** Phylogenetic generalized least-squares analyses of absolute transposable element abundance.

**Supplementary Table S12.** Curated FitMG94-derived *dN/dS* dataset for high-confidence phylogenetic branches.

